# Down Regulation Of IL-1β Secretion By Tgf-β1 In Macrophages Infected With Dengue Virus

**DOI:** 10.1101/2021.02.03.429556

**Authors:** Brenda Ramírez-Aguero, Javier Serrato-Salas, José Luis Montiel-Hernández, Judith González-Christen

## Abstract

Several pathogenic mechanisms have been linked to the severity of dengue virus infection, like viral cytotoxicity, underlying host genetics and comorbidities such as diabetes and dyslipidemia. It has been observed that patients with severe manifestations develop an uncontrolled immune response, with an increase in pro-inflammatory cytokines such as TNF, IL-1β, IL-8, IL-6 and chemokines that damage the human microvascular endothelium, and also in anti-inflammatory cytokines IL-4, IL-10 and TGF-β1. The role of TGF-β1 on dengue is not clear; few studies have been published, and most of them from patient sera data, with both protective and pathological roles have described. The aim of this study was to evaluate the ability of TGF-β1 to regulate the secretion of IL-1β in macrophages infected by DENV using THP-1 cells treated with recombinant TGF-β1 before or after DENV infection. By RT-PCR we did not observe a difference in IL-1β expression between infected cells pretreated with TGF-β1 and those that were not. However, secretion of IL-1β was reduced only in cells stimulated with TGF-β1 before infection, and not in those treated 2 hours post-infection. TGF-β1 receptor blockage with SB505124 inhibitor, prior to the addition of TGF-β1 and infection, abrogated the inhibitory effect of TGF-β1. Our results suggest that DENV could regulate the function of TGF-β1 on macrophages. This negative regulation of the TGF-β1 pathway could be used by DENV to evade the immune response and could contribute to the immunopathology.

## Introduction

Despite health prevention campaigns and the recent application of a licensed vaccine, mortality and the disease burden metrics, such as years of life lost (YLLs), associated to Dengue increased by 65.5% and 32% respectively, from 1998 to 2017(Collaborators 2018).

Humans infected by dengue virus (DENV) can present a wide array of manifestations but the great majority will not develop a serious conditions, 20 to 25% will be symptomatic and only a small proportion will develop clinically relevant complications, including secondary symptoms associated to capillary leakage and organ involvement that will require hospitalization (Martinez, Garza et al. 2019, Wilder-Smith, Ooi et al. 2019). Patients with dengue have a general increase in cytokine titers and severity has been associated to at least 25 different serum molecules, mainly TNF, IFNγ, IP-10, RANTES, chemokines, IL-1β, IL-4, IL-6, IL-7, IL-8, IL-10 and TGF-β1(Pandey, Jain et al. 2015, Soo, Khalid et al. 2017, Srikiatkhachorn, Mathew et al. 2017, Patra, Mallik et al. 2019). It has been proposed that the disease is primarily immune mediated, because the dysregulation of TNF and IL-6, among others, leads to vascular alterations, plasma leakage, thrombocytopenia and organ damage (St John and Rathore 2019).

IL-1β is a prototype of inflammatory cytokine that has been associated to diverse pathologies, usually in combination with other cytokines such as TNF or IL-6. The role of IL-1β in dengue pathogenesis is not completely understood. Studies in humans have shown contradictory results, some associated with an increase in the development of hemorrhagic symptoms (or severity of dengue), while others do not find a relationship at all (Chaturvedi, Nagar et al. 2006, Bozza, Cruz et al. 2008).

As in the case of IL-1β, the role of TGF-β1 on dengue pathogenesis is not clear. Some studies in patients associated an increase in TGF-β1 with severity because levels were reduced in controls or in patients with dengue fever compared to dengue hemorrhagic fever; this difference seems to be more important particularly the day before the drop in temperature (Laur, Murgue et al. 1998, Agarwal, Elbishbishi et al. 1999, Patra, Mallik et al. 2019). However, polymorphism studies in dengue patients associated the genotypes of high TGF-β1 producers with controls (Perez, Sierra et al. 2010, Yeo, Azhar et al. 2014). In addition, Tillu et al found a greater number of platelets in those patients with higher concentration of Tregs and TGF-β1 (Tillu, Tripathy et al. 2016), suggesting a protective role for TGF-β1.

*In vitro*, different studies have observed a relationship between TGF-β and IL-1β. It has been observed that Smad6/7 proteins produced in response to TGF-β signaling induce a negative regulation of IL-1R/TLR signaling (Lee, Kim et al. 2010). Also, TGF-β can inhibit the production of IL-1β in response to LPS through downregulation of CD14 (Imai, Takeshita et al. 2000). However, the relationship between these cytokines in the context of clinically expressed Dengue infection is still unknown. Therefore, we evaluated the role of TGF-β1 in the regulation of IL-1β production by macrophages infected with DENV.

## Materials and Methods

### Virus, cell culture, differentiation and infection of THP-1 cells

Dengue Virus serotype 2 New Guinea C was kindly donated by Dr. Humberto Lanz Mendoza, stock aliquots of 20 μl were stored at −70°C, and diluted in Dubelcco’s-PBS just before infection (SIGMA-Aldrich). THP-1 cells (ATCC Tib 202) were grown in Advance-RPMI 1640, supplemented with 3% Fetal Bovine Serum, (complete medium; all from GIBCO-Thermo Scientific) at 37°C and 5% CO_2_. For differentiation into macrophages, cells were treated with 10 nM PMA (SIGMA-Aldrich) for 72 hours before infection or stimulation. Cells were differentiated in 24 well cell culture plates (Corning), 2×10^5^ cell/well, with sterile round glass coverslips for microscopy or ELISA quantification, or in 25 cm^2^ flasks (Corning) at 4×10^6^ in 10 mL for RT-PCR assays. For infection, cells were washed with PBS, Dengue virus was added at a MOI=1 in a minimum volume of Dubelcco’s PBS with CaCl_2_ and antibiotics (SIGMA-Aldrich). After 2 hours, cells were washed with PBS and were further incubated for 24 hours in fresh complete medium. Infection was confirmed by fluorescent microscopy using anti-Dengue 1:20 (ab155042, AbCAM), anti-mouse Alexa 488 1:200 (Jackson Immunology), or by RT-PCR for prM/M-C region.

### Effect of TGF-β1 prior to infection with DENV on IL-1β expression and secretion

For assays of effects of TGF-β1 on the expression and secretion of IL-1β, differentiated cells were washed with PBS and fasted for 2 hours in RPMI-1640 (Gibco, Thermo Scientific) without serum, then 20 ng/mL of recombinant TGF-β (PepProtech Inc.) was added and incubated for an additional 3 hours. Cells were washed in PBS and infection proceeded as described.

### Effect of TGF-β1 post infection with DENV over IL-1β expression and secretion

For the post-infection assay, fasted cells were infected with DENV in PBS for 2 hours as described, and then complete medium was added with 20 ng/mL of TGF-β1 (PepProtech Inc.). Cells were further incubated for an additional 24 hours.

### Effect of SB 505124 on IL-1β secretion

Differentiated THP-1 cells were washed with PBS, fed with RPMI-1640 without serum, containing 1 mM of SB505124 inhibitor (TOCRIS Biosciences) and incubated for 2 hours. Then, 20 ng/mL of TGF-β1 was added, incubated for 3 hours before infection as described previously.

### Stimulation of THP-1 with LPS

LPS was used to induce an inflammatory profile on THP-1 differentiated with PMA. The cells were processed as for DENV infection, but in the final feeding with fresh complete medium, 10 ng/mL of LPS from *E. coli* O11:B4 was added (Sigma-Aldrich).

### Quantification of secreted IL-1β

In all assays, 24 hours after infection, the supernatant was withdrawn, centrifuged at 575g for 10 min (Spectrafuge, Labnet), aliquoted and stored at −20°C until used. In order to measure secreted IL-1β we used the commercial Human IL-1β ELISA MAX kit (Biolegend), according to the manufacturer’s instruction.

### Total RNA isolation and reverse transcription-polymerase chain reaction (RT-PCR)

Total RNA was isolated from THP-1 cell lysate in Tri-Reagent (SIGMA-Aldrich) using a standard organic solvents precipitation protocol according to the manufacturer’s instruction. Concentration/quality of total RNA was measured at 280/260 nm in a microvolume detection Take3 plate in an Epoch microplate spectrophotometer (Bioteck), and agarose gel electrophoresis. Complementary DNA (cDNA) was then synthetized from 1 μg of total RNA using the Revert Aid First Strand cDNA Synthesis Kit (Thermo Scientific), employing random hexamer primers. PCR was performed using PCR Master Mix (2x) (Thermo Scientific) in a 3PRIME Thermal Cycler (TECHNE, Bibby Scientific), according to manufacturer’s instructions. Reaction conditions were, 35 cycles of denaturalization at 95°C for 30 sec, annealing at specific temperature for each pair of primer, 60 sec and extension 72°C for 30 sec. Specific primers and annealing temperature in °C for IL-1β (57°C) forward 5’GTCATTCGCTCCCACATTCT3’, reverse 5’ACTTCTTGCCCCCTTTGAAT3’;(Bozza, Cruz et al. 2008); β actin (57-60°C) forward 5’ AGGATCTTCATGAGGTAGT3’, reverse 5’GCTCCGGCATGTGCAA3’, TGF-β1 (60°C) forward 5’GGTACCTGAACCCGTGTTGCT3’, reverse 5’TGTTGCTGTATTTCTGGTACAGCTC3’. Products were analyzed by agarose gel electrophoresis in a ChemiDoc XRS-Quantity One Image Analyzer (BioRad)

### DENV RNA genome detected by PCR

A PCR assay was performed to detect infection and to amplify the RNA strand of DENV. Total RNA of infected cells and synthesis of cDNA was conducted as described for IL-1β. cDNA reaction was adjusted to 500 ng/μl and used for PCR in Master Mix 2x (Thermo Scientific), using a 3PRIME Thermal Cycler (TECHNE, Bibby Scientific). Reaction conditions were: 35 cycles denaturalization at 95°C for 20 sec, annealing at 60°C for 30 sec and extension at 72°C for 60 sec. The first PCR included the following primers DENV ALL RV 5’CCCCATCTATTCAGAATCCCTGC3’ and DENV ALL F 5’CAATATGCTGAAACGAGAGAGAA3'. A nested PCR was performed with the amplicon from first PCR product diluted 1:100 using the same reaction conditions for cycling, with the following pair of oligonucleotides DENV ALL F 5’CAATATGCTGAAACGAGAGAGAA3 and DENV2RV 5’TGCTGTTGGTGGGATTGTTA 3’.

### Statistical analysis

Descriptive statistics was used employing media and standard deviation (SD). Comparison between experimental conditions was performed by using unpaired Students t-tests and p<0.05 values were considered as statistically significant. Graphs and statistic were performed using Prism v.8.1.0 software (GraphPad).

## Results

### Treatment with TGF-β1 reduces IL-1β secretion only if it is applied before infection

In our experimental conditions, with a MOI=1 at 24 hours after infection, we observed 40% of infected cells, confirmed by fluorescent microscopy and RT-PCR (Supplementary material). In these conditions, infection with DENV induces the secretion of IL-1β (p<0.005). This stimulus was lower than that induced by LPS. In order to determine if TGF-β1 could have a regulatory effect on IL-1β secretion, we treated THP-1 differentiated cells with 20 ng/mL of recombinant TGF-β1 for 3 hours previous infections. This treatment resulted in a significant inhibition of the IL-1β secretion (p<0.03) only in infected cells, but not in cells treated with LPS (Figure 1A). However, this effect was no longer sustained if cells were first infected and then 2 hours later, treated with recombinant TGF-β1 (p=0.46), (Figure 1B).

**Figure 1.**
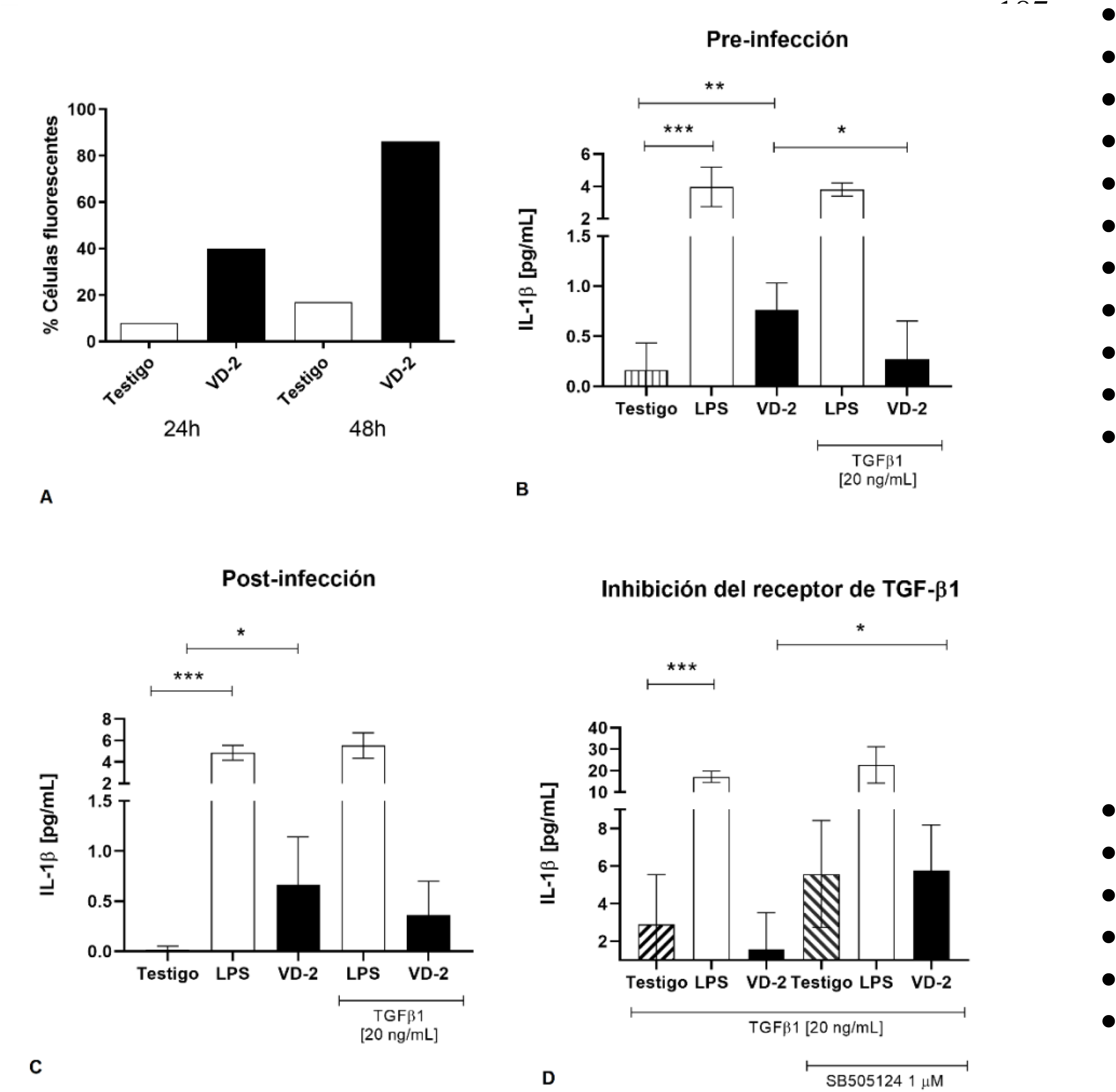
A. Percentage of fluorescent cells. Controls at 24 and 48 hpi, average 10-15% positive cells. DENV 24 and 48 hpi. 40% and 85% positive cells. B. IL-1β secretion in experimental groups adding TGF-β1 recombinant previous to infection. C. IL-1β secretion in cells adding TGF-β1 before viral infection. D. IL-1β secretion in different treatments using TGF-β1 receptor competitive inhibitor. Statistical significative differences were made using Student’s T assay. P<0.05 *, p<0.01 **, p<0.005 ***.

**Figure 2.**
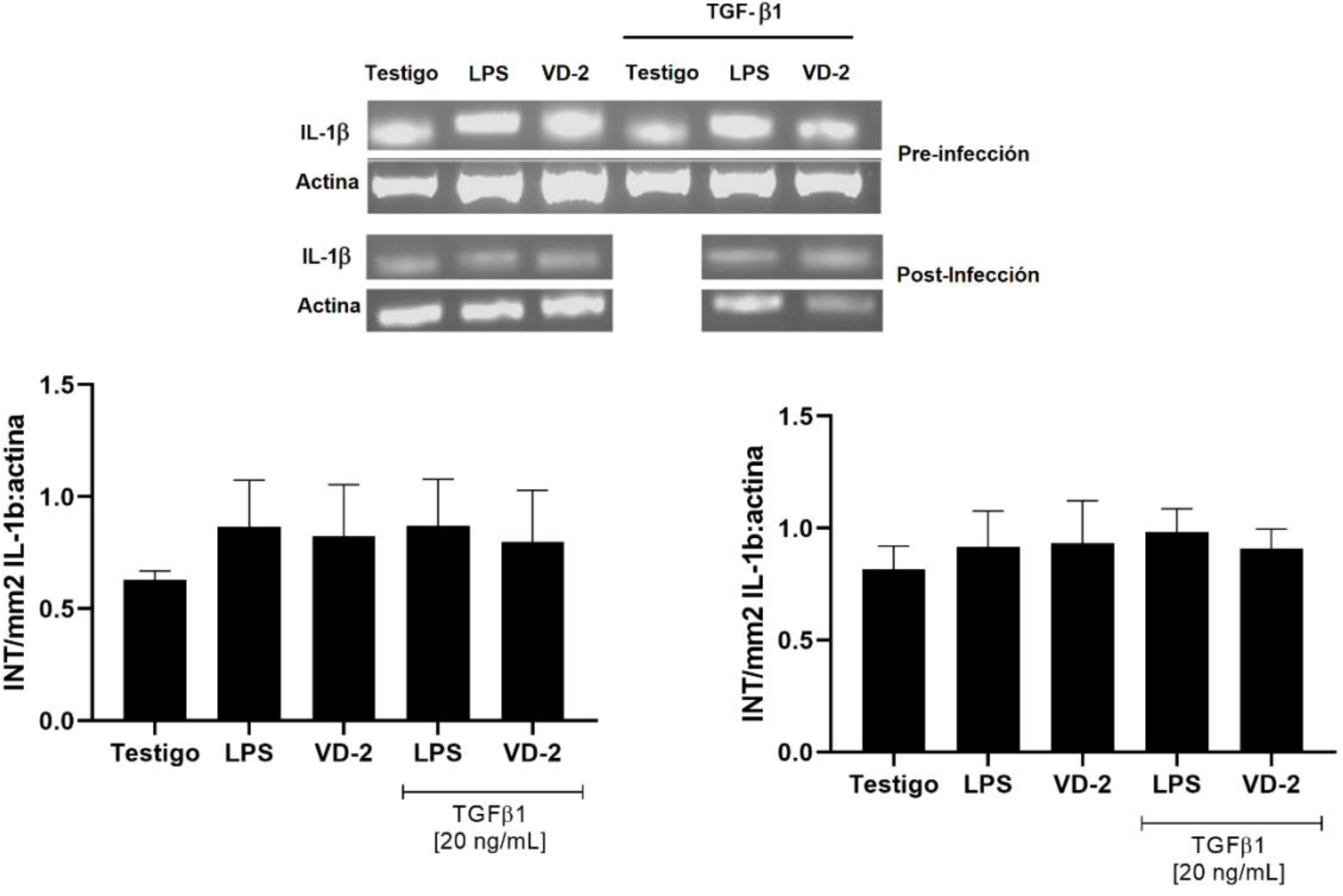
Previous treatment with TGF-β1 does not modify mRNA IL-1β. A. mRNA IL-1β and β-actin in cells with TGF-β recombinant or PBS over 3 h before and after viral infection. B. Relative expression of IL-1β on TGF-β1 previous to infection was normalized to actin as pixel intensity per mm^2^. C. Relative expression of IL-1β on TGF-β1 added post-infection was normalize to actin as pixel intensity per mm^2^

### Inhibition of TGF-β1 receptor abrogates the effect of TGF-β1 1 over IL-1β

In order to further validate this data, pretreatment of the THP1 with a selective inhibitor of TGF-βRI, SB505124, was performed prior to exposure to 20 ng/mL of TGF-β1; 3 hours later infection with DENV proceeded as described above. This treatment abrogated the inhibitory effect of TGF-β1 on infected cells. In fact, after inhibitory pretreatment, we observed a higher increase of IL-1β secretion induced by DENV infection (p<0.04), in comparison to non-inhibited cells. It is noteworthy that both the uninfected cells and those treated with LPS also responded with an increase in the production of IL-1β (Figure 1C).

### Effect of TGF-Β1 over IL-1β secretion is not related with IL-1β expression

IL-1β is produced as a 31 kDa precursor that needs to be processed for its secretion. In order to determine if the effect of TGF-β1 is due to a repression of IL-1β expression, we evaluated the IL-1β messenger using RT-PCR. Although infection and LPS treatment increased the expression of IL-1β RNA messenger, we did not observe any change when comparing the infected cells without treatment with those who underwent treatment with TGF-Β1, suggesting that the effect of TGF-β1 is major associated to secretion.

## Discussion

The complexity of the disease caused by the infection of DENV is due to the fact that it depend on many factors, such as virus serotype and genotype, the mosquito proteins, previous infection with dengue or other arbovirus, host genetic polymorphism, co-morbidities such as obesity, cardiovascular disease or asthma (St John and Rathore 2019). However it has been observed that the immune response of the host plays an important role in the pathogenesis of clinically expressed Dengue because it leads to an increase in inflammatory and anti-inflammatory cytokines (Chaturvedi, Nagar et al. 2006, Pang, Cardosa et al. 2007, St John and Rathore 2019) There is no clear proposal that explains how this cytokine imbalance can lead to a worsening of the disease, but one hypothesis is that different cytokines released during infection, such as TNF, IL-6 or IL-8, has a direct action on vasculature, leading to dysregulation of the endothelial cell barrier associated to increased vascular permeability, resulting in pleural effusion or ascites (Srikiatkhachorn, Mathew et al. 2017). The search of biomarkers that can predict the severity of disease before symptoms appears has been going on for years. The 24 to 48 hour period before the drop in body temperature is especially important because vascular alterations, plasma leakage, increase hematocrit, lowering of platelets and organ commitment usually occurs 2-3 days after deffervescence (Yacoub and Wills 2014). So far it has not been possible to determine these biomarkers, not only because of patient variability, but because of methodological differences that include disease severity classifications, age of patients, the day on which the serum is obtained, etc (John, Lin et al. 2015).

At the end of 90’s, several clinical studies associated the increased levels of TGF-β1 with severity because levels were reduced in controls or in patients with dengue fever compared to dengue hemorrhagic fever; this difference seems to be more important particularly the day before the drop in temperature, and last for 9 day after fever onset (Letterio and Roberts 1998, Agarwal, Elbishbishi et al. 1999, Patra, Mallik et al. 2019). However, polymorphism studies comparing healthy or asymptomatic patients with those that developed severe dengue associated the genotypes of high TGF-β1 producers with controls, suggesting a protective role for TGF-β1 (Perez, Sierra et al. 2010, Yeo, Azhar et al. 2014). Interestingly Djamiatun et al. found that TGF-β1 plasma levels always were lower than healthy children, even if there was an increase in patients with hemorrhagic manifestations (Djamiatun, Faradz et al. 2011). Otherwise *in vitro* study has shown that cells from a DENV serotype donor, produces higher TGF-β1 when re-challenged with the same serotype, but not with a heterologous serotypes. As severe dengue has been associated to reinfection, this study could also suggest a protective role for TGF-β1(Perez, Sierra et al. 2010, Sierra, Perez et al. 2010)

TGF-β1 is a multifunctional cytokine which immunomodulatory profile changes depending on concentration, tissue location and context of action; for example on T cells, leading to the suppression of Th1, Th2 and CD8 activation and also favoring the generation of FoxP3/Tregs (Li and Flavell 2006, Chen and Ten Dijke 2016). Tillu et al. found elevated Tregs frequencies in patients with mild dengue disease, suggesting a protective role of these cells, and also noted a relationship between TGF-β1 levels and greater number of platelets in those patients with higher concentration of Tregs (Tillu, Tripathy et al. 2016). On the other hand, effects of TGF-β1 on macrophages can be either stimulating or inhibitory, depending on the presence of other cytokines and the state of maturation (Letterio and Roberts 1998, Travis and Sheppard 2014)

In order to establish if there is an association between TGF-β1 and the macrophage response to DENV infection, we conducted an *in vitro* study using THP1 differentiated into macrophages. In our assay’s condition, at 24 hours post-infection, 40% of cells were positive for DENV but we observed a significant increase in IL-1β secretion. Other groups also have observed that macrophages infected with DENV, even in the absence of antibodies, secrete IL-1β, up until 6 hours post-infection (Callaway, Smith et al. 2015). Here we observed a total reduction of IL-1β secretion in infected cells that were treated 3 hours with TGF-β before infection, but not in cells stimulated with LPS. Concentration of IL-1β released by macrophages stimulated by LPS was almost 5 times higher than those infected by DENV. This difference could be explained by the fact that LPS and DENV bind to different TLRs.

In our assay of pre-treated THP-1 cells with a selective inhibitor of TGF-βI receptor, SB-505124, that inhibit specific activation dependent of Smad2/3, the inhibitory effect of the TGF-β on the secretion of IL-1β was abrogated. Additionally, once the infection started, we no longer observed the inhibitory effect of TGF-β1 on IL-1β secretion. For other part, we did not notice any change in IL-1β RNA messenger using RT-PCR, suggesting that the mechanism of TGF-β1 should be related to events after the expression of IL-1β; however, it is necessary to confirm this data by qPCR. Kwon et al., using siRNA assay shown that knock-down of Smad7, an inhibitory protein of TGF-β signaling, is associated to a lesser infection. They also showed that infected U937-DC-Sign cells increased the expression of Smad7 in response to DENV (Kwon, Heo et al. 2014). At this respect, we can assume that DENV should manipulate the response to TGF-β1 potentially by modification of the cellular pathways that is why it should be important to evaluate the changes in expression and activation of Smad 2/3 and Smad7.

IL-1β is produced as a 34 kDa precursor that requires maturation by proteases, usually Caspase-1, before secretion. IL-1β is a leaderless cytosolic protein and the precise mechanism of release of this cytokine is not completely understood, but it is known that it requires a non-canonical pathway independent of Golgi. Several mechanisms have been proposed to explain the release of this cytokine as pores associated to phosphatidylserine, autophagy vesicles or pyroptosis (Wu, Chen et al. 2013, Brough, Pelegrin et al. 2017, New and Thomas 2019).

We did not find previous evidence that TGF-β1 has an effect on the inhibition of secretion of IL-1β that could be associated to the activation of caspase-1 and induction of pyroptosis. Some studies have shown that DENV infected monocytes or macrophages died by pyroptosis after activation of caspase-1 (Tan and Chu 2013, Wu, Chen et al. 2013). Another study has shown the role of caspase-4 on the activation the maturation of IL-1β in DENV infected cells (Cheung, Sze et al. 2018). Interestingly, the secretion of IL-1β was observed almost 90 hours before the activation of caspase-1 or cell death. It is known that IL-1β could be processed by other proteases and not necessarily due to its secretion leading to cell death (Netea, van de Veerdonk et al. 2015). In contrast, here we observed elevated IL-1β secretion at 24 hours post-infection as others have reported (Callaway, Smith et al. 2015).

Autophagy has a dual role in the regulation of IL-1β because it is required both for the maturation and release, and also controls its release by its degradation before secretion. The autophagy vesicle required for maturation and release of IL-1β seems to be a non-classical vesicle. IL-1β is associated to TRIM16 and after complex traffic to an intermediate membrane positive for LC3-II, and then a vesicle is formed and associated to SEC22b. In order to reach the plasma membrane this vesicle undergoes a SNARE-modified fusion (New and Thomas 2019). However, inhibitors of autophagy, or genetic defective autophagy cells, result in an increase of IL-1β secretion, but not of TNF or IL-6 on cells stimulated with LPS. The autophagy vesicle formed in absence of an inflammasome-inducing signal is independent of TRIM16 and co-localize with LAMP-1 and STX17, leading to transport and degradation of IL-1β by lysosomes (Harris 2011, Harris 2013).

During viral infection, the virus manipulates different cellular pathways, such as the response to IFNs and autophagy. Several studies have shown that many autophagy induced pathways are activated by DENV for their replication (Datan, Roy et al. 2016). Inhibition of ER stress signaling limits the ability of DENV to induce autophagy and decrease the formation of mature virions (Datan, Roy et al. 2016). It has been shown that proteins of the viral replication complex do not co-localize with autophagosome membranes, suggesting that it may play another role, such as immune evasion response (Heaton and Randall 2011). On the other hand, several studies have shown that TGF-β1 increases autophagy, mediated through the Smad and JKN pathways.

Our proposal is that in the absence of DENV infection, TGF-β1 will induce the accumulation of IL-1β and inflammasome proteins in autophagic vesicles associated with STX17 and SNARE, directing the complex to lysosomes. But when the dengue virus initiates the infection of macrophages, mediated by TLR 3,4 and 7, the signal pathways lead to the accumulation of IL-1β in vesicles associated to TRIM16 and SEC22b, eventually resulting in the secretion of IL-1β. Evidently this proposal must be confirmed by experimental data. Another mechanism that could explain why infected DENV macrophages do not respond to TGF-β1 could be associated to signaling downstream pathways.

A limitation of our study is that we performed our assays in the absence of antibodies. It is known that the macrophage activation pathways are very different if the virus binds directly to the macrophages or is mediated by immunocomplexes and FcRγ. Callaway et al. have shown that Dengue-Antibody complexes can increase the secretion of IL-1β, but only when the immunocomplex is generated by serum homologous to the serotype of the virus and not with heterologous sera (Callaway, Smith et al. 2015). This is also important because it is known that TGF-β1 reduces the expression of FcRγ on the surface of cells.

In conclusion, this study supports the proposal that TGF-β1 may have a limiting role in the development and severity of dengue disease, but only on those individuals who prior to infection have TGF-β present in the cellular environment.

## Author Contributions

RB, MJ and GJ contributed to conception, design and interpretation of results of the study. RB conducted the main laboratory work. SJ conducted the test for PCR detection of DENV and wrote material and methods section. All authors contributed to read, write and approval the final manuscript.

## Acknowledgments

We thank PhD Humberto Cuahutemoc Lanz Mendoza, form CISEI-INSM for his support in carrying out some experiments in his laboratory, as well as the donation of viral stock. PhD Daniel Xibille Fredman by careful reading and correction of the text.

## Data Availability Statement

The dataset and raw data supporting the conclusion of this manuscript will be made available by authors, without undue reservation to any qualified researcher.

---

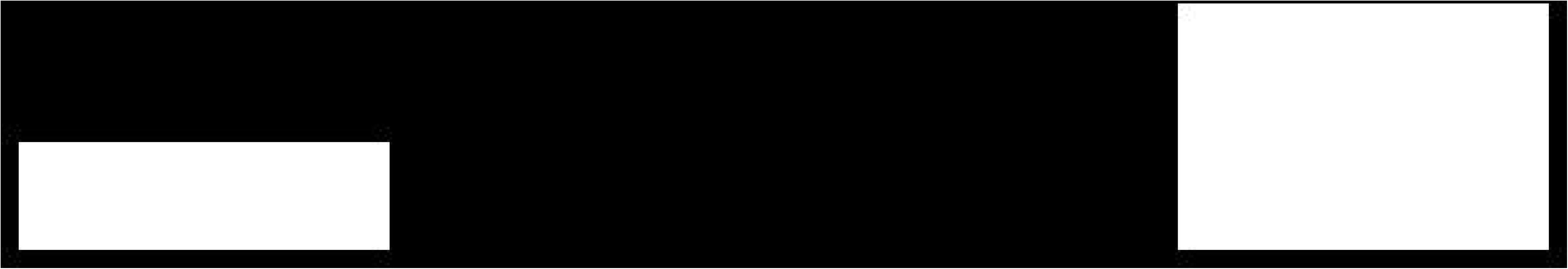

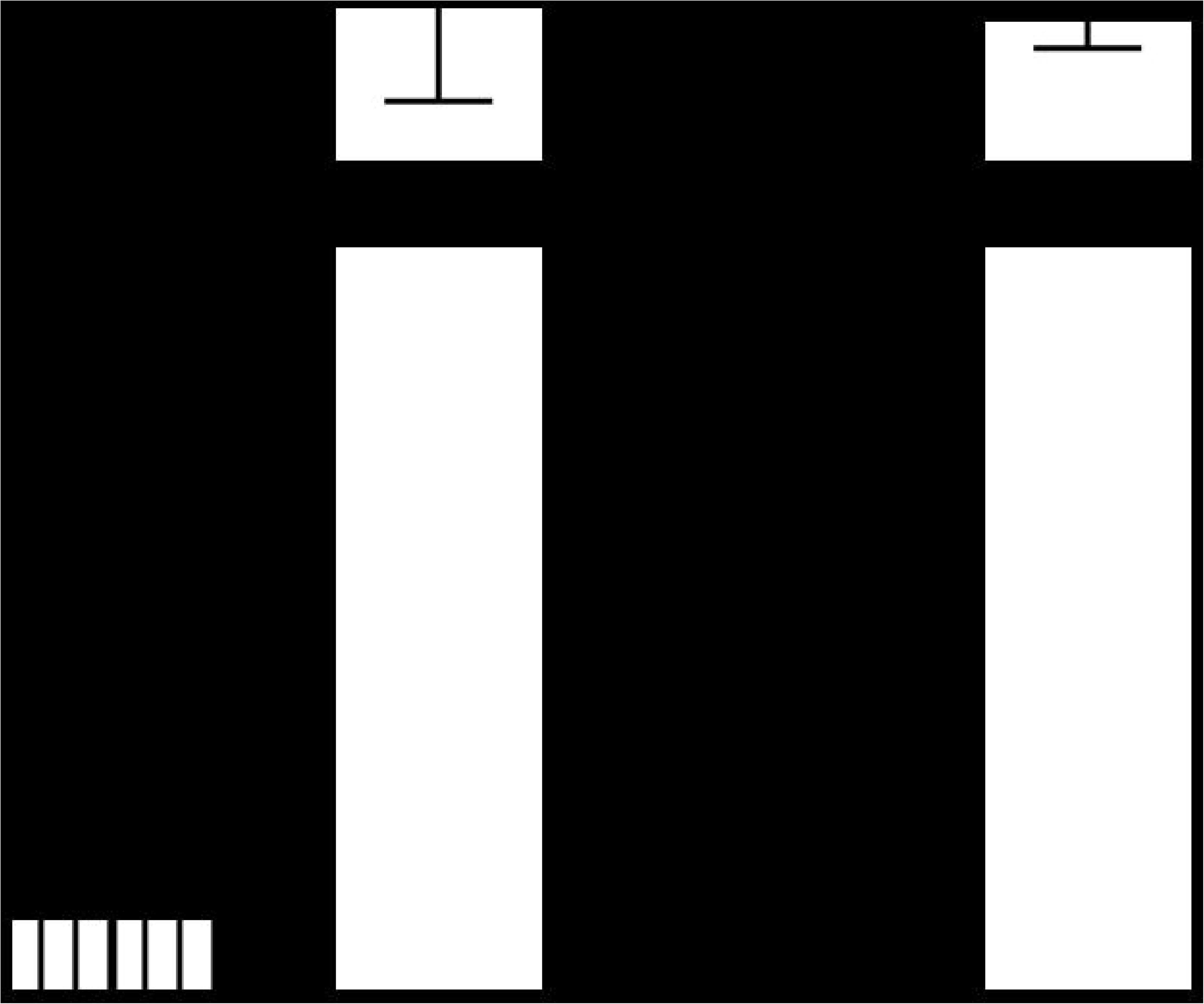

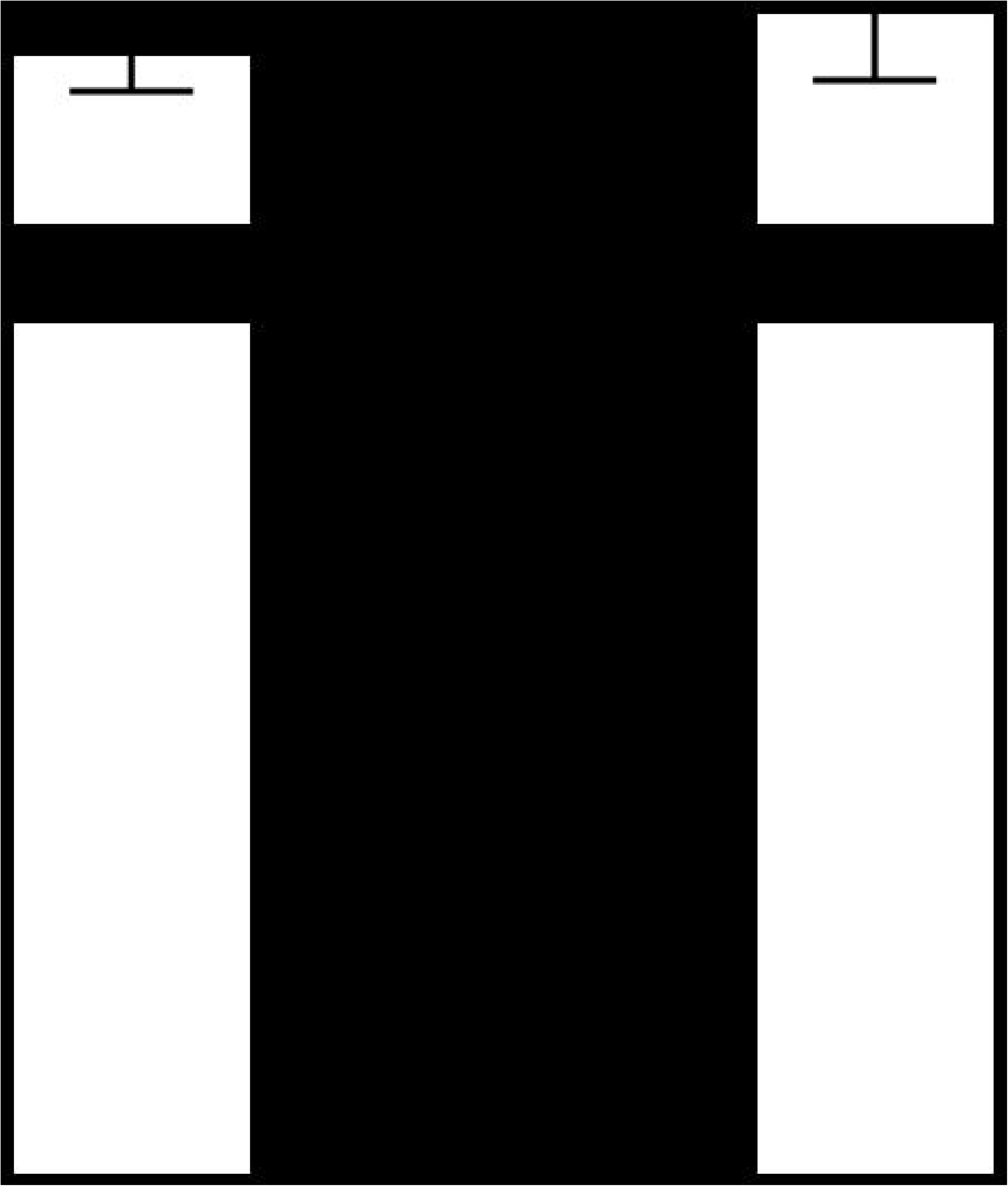

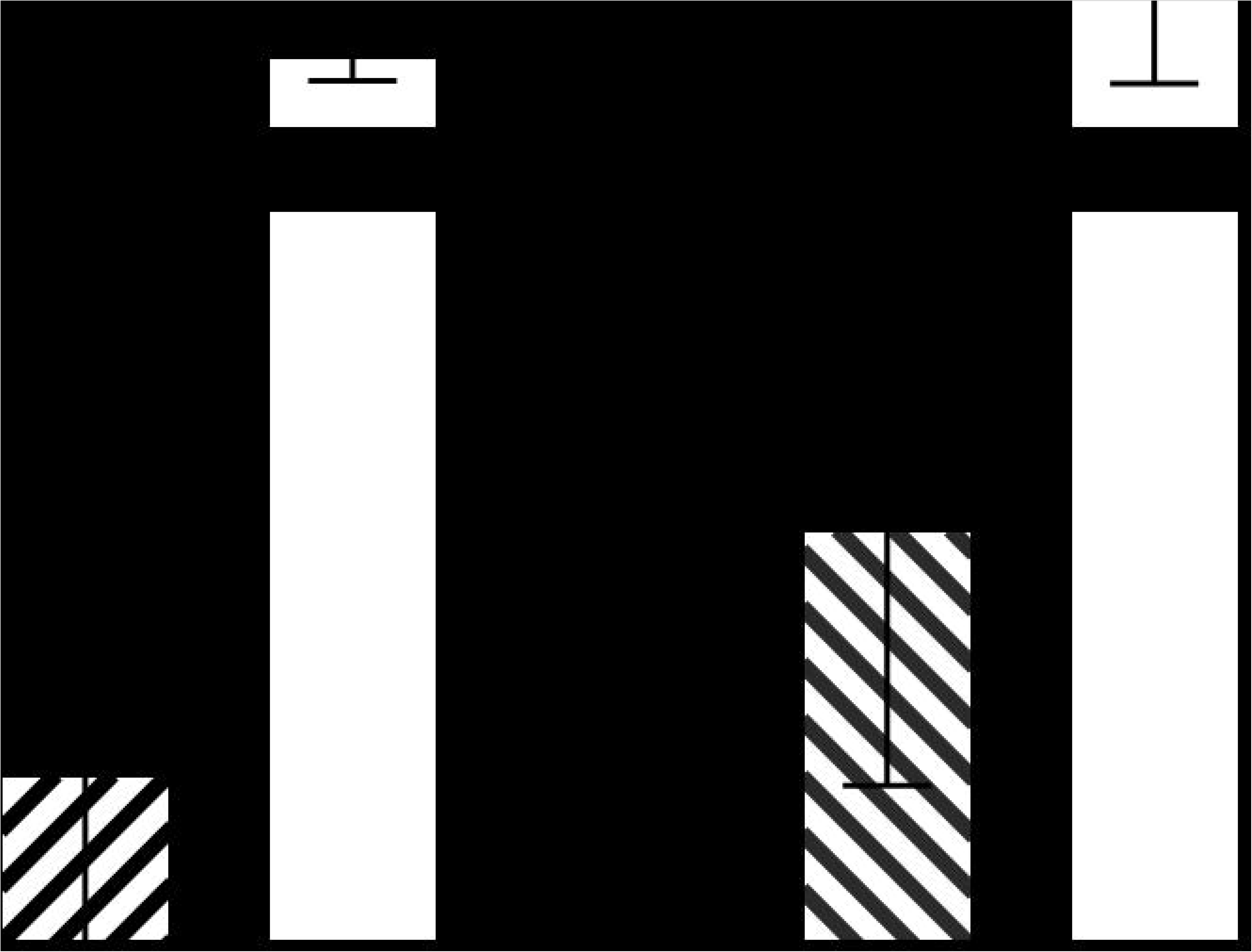

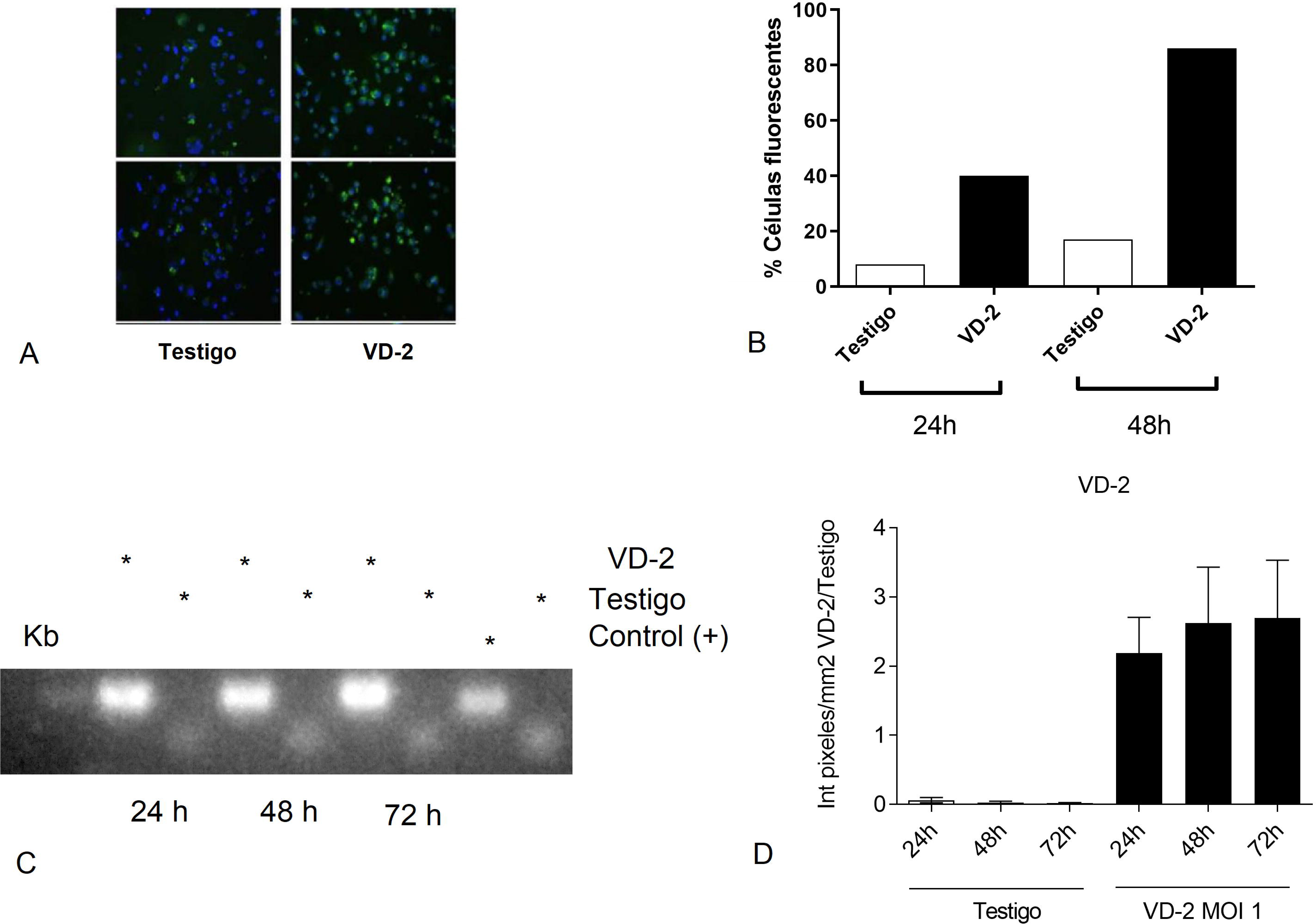

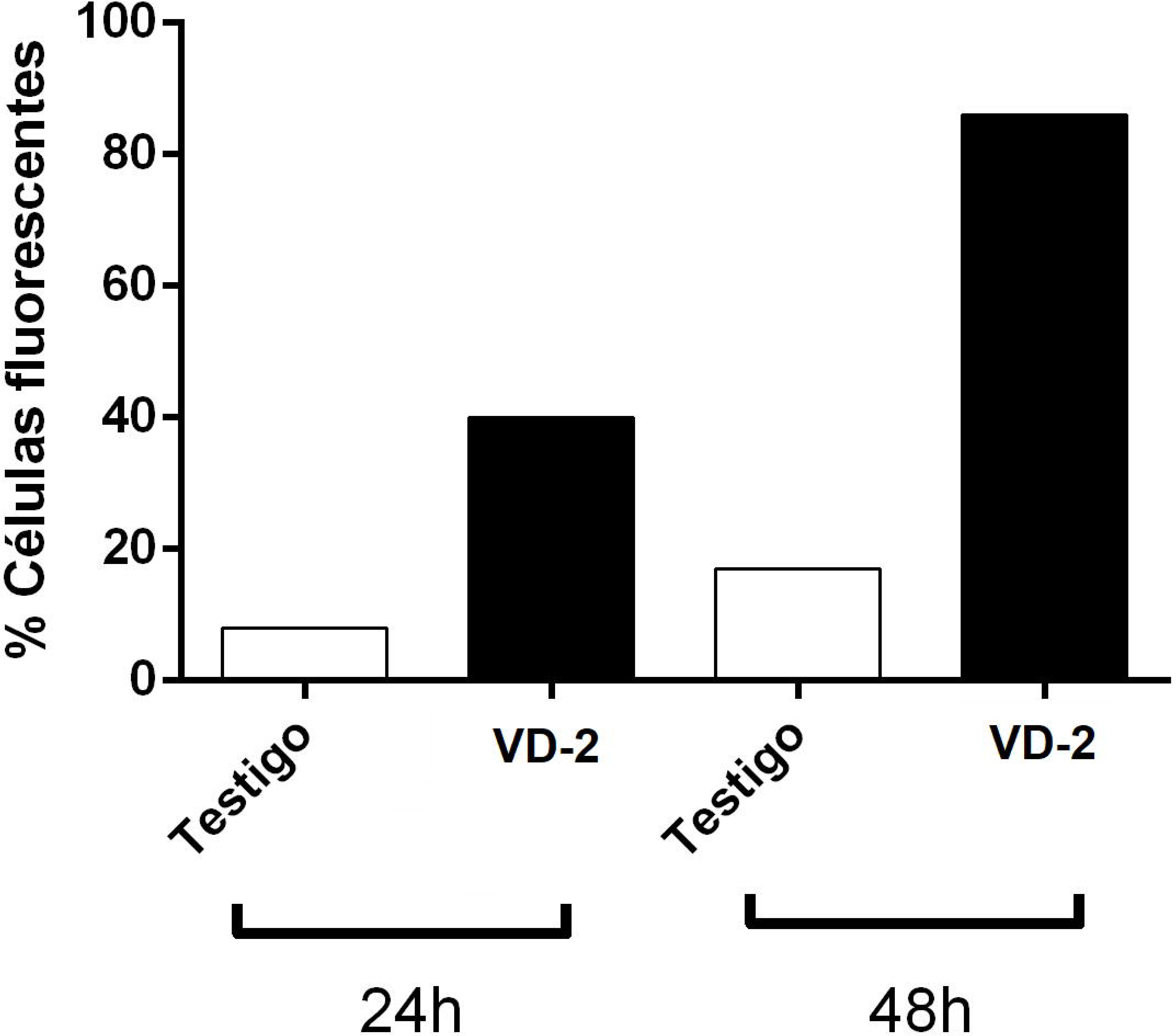

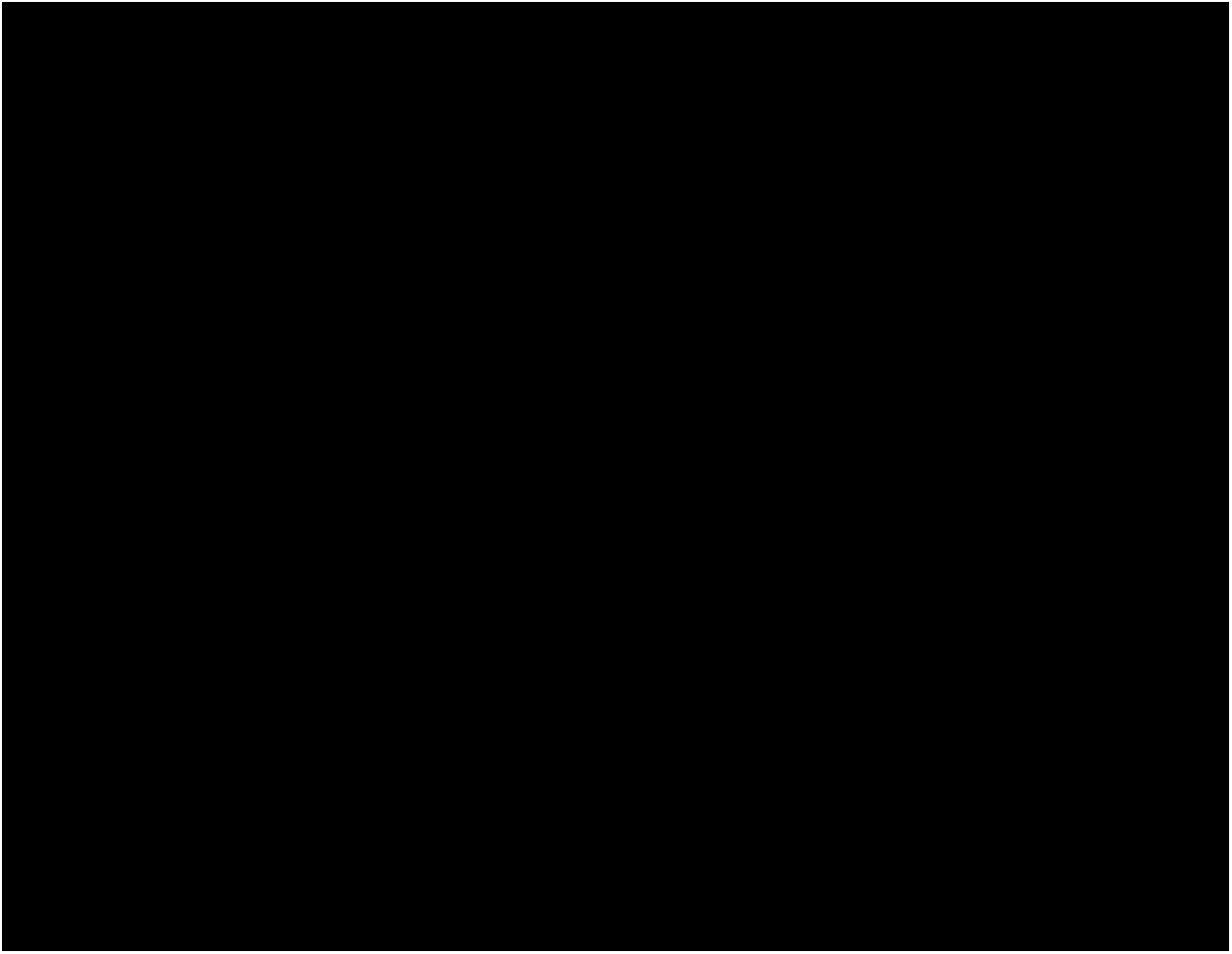

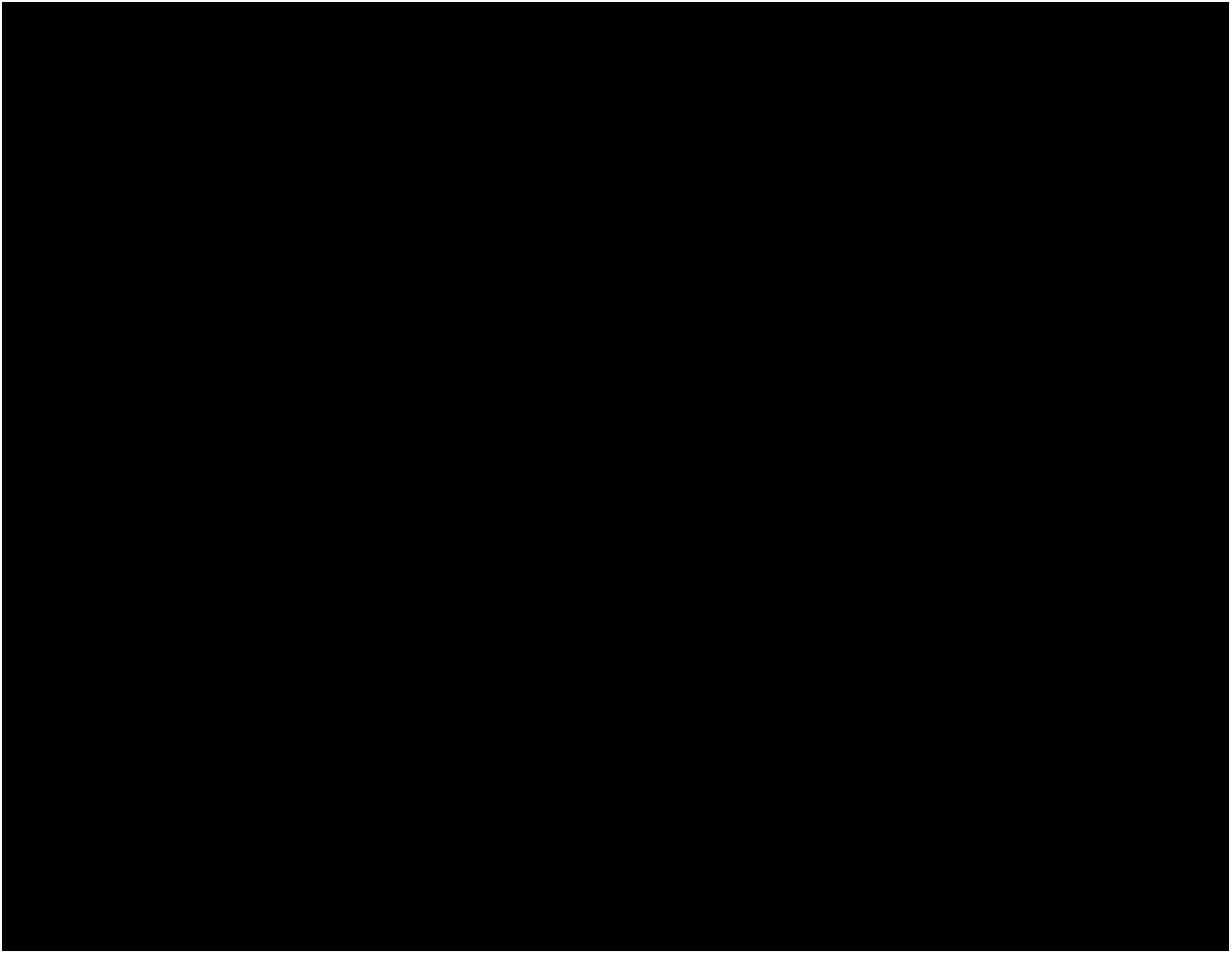

## References

Agarwal, R., E. A. Elbishbishi, U. C. Chaturvedi, R. Nagar and A. S. Mustafa (1999). “Profile of transforming growth factor-beta 1 in patients with dengue haemorrhagic fever.” Int J Exp Pathol 80(3): 143–149.

Bozza, F. A., O. G. Cruz, S. M. Zagne, E. L. Azeredo, R. M. Nogueira, E. F. Assis, P. T. Bozza and C. F. Kubelka (2008). “Multiplex cytokine profile from dengue patients: MIP-1beta and IFN-gamma as predictive factors for severity.” BMC Infect Dis 8: 86.

Brough, D., P. Pelegrin and W. Nickel (2017). “An emerging case for membrane pore formation as a common mechanism for the unconventional secretion of FGF2 and IL-1beta.” J Cell Sci 130(19): 3197–3202.

Callaway, J. B., S. A. Smith, K. P. McKinnon, A. M. de Silva, J. E. Crowe, Jr. and J. P. Ting (2015). “Spleen Tyrosine Kinase (Syk) Mediates IL-1beta Induction by Primary Human Monocytes during Antibody-enhanced Dengue Virus Infection.” J Biol Chem 290(28): 17306–17320.

Chaturvedi, U. C., R. Nagar and R. Shrivastava (2006). “Macrophage and dengue virus: friend or foe?” Indian J Med Res 124(1): 23–40.

Chen, W. and P. Ten Dijke (2016). “Immunoregulation by members of the TGFbeta superfamily.” Nat Rev Immunol 16(12): 723–740.

Cheung, K. T., D. M. Sze, K. H. Chan and P. H. Leung (2018). “Involvement of caspase-4 in IL-1 beta production and pyroptosis in human macrophages during dengue virus infection.” Immunobiology 223(4-5): 356–364.

Collaborators, G. B. D. C. o. D. (2018). “Global, regional, and national age-sex-specific mortality for 282 causes of death in 195 countries and territories, 1980-2017: a systematic analysis for the Global Burden of Disease Study 2017.” Lancet 392(10159): 1736–1788.

Datan, E., S. G. Roy, G. Germain, N. Zali, J. E. McLean, G. Golshan, S. Harbajan, R. A. Lockshin and Z. Zakeri (2016). “Dengue-induced autophagy, virus replication and protection from cell death require ER stress (PERK) pathway activation.” Cell Death Dis 7: e2127.

Djamiatun, K., S. M. Faradz, T. E. Setiati, M. G. Netea, A. J. van der Ven and W. M. Dolmans (2011). “Increase of plasminogen activator inhibitor-1 and decrease of transforming growth factor-b1 in children with dengue haemorrhagic fever in Indonesia.” J Trop Pediatr 57(6): 424–432.

Harris, J. (2011). “Autophagy and cytokines.” Cytokine 56(2): 140–144.

Harris, J. (2013). “Autophagy and IL-1 Family Cytokines.” Front Immunol 4: 83.

Heaton, N. S. and G. Randall (2011). “Dengue virus and autophagy.” Viruses 3(8): 1332–1341.

Imai, K., A. Takeshita and S. Hanazawa (2000). “Transforming growth factor-beta inhibits lipopolysaccharide-stimulated expression of inflammatory cytokines in mouse macrophages through downregulation of activation protein 1 and CD14 receptor expression.” Infect Immun 68(5): 2418–2423.

John, D. V., Y. S. Lin and G. C. Perng (2015). “Biomarkers of severe dengue disease - a review.” J Biomed Sci 22: 83.

Kwon, Y. J., J. Heo, H. E. Wong, D. J. Cruz, S. Velumani, C. T. da Silva, A. L. Mosimann, C. N. Duarte Dos Santos, L. H. Freitas-Junior and K. Fink (2014). “Kinome siRNA screen identifies novel cell-type specific dengue host target genes.” Antiviral Res 110: 20–30.

Laur, F., B. Murgue, X. Deparis, C. Roche, O. Cassar and E. Chungue (1998). “Plasma levels of tumour necrosis factor alpha and transforming growth factor beta-1 in children with dengue 2 virus infection in French Polynesia.” Trans R Soc Trop Med Hyg 92(6): 654–656.

Lee, Y. S., J. H. Kim, S. T. Kim, J. Y. Kwon, S. Hong, S. J. Kim and S. H. Park (2010). “Smad7 and Smad6 bind to discrete regions of Pellino-1 via their MH2 domains to mediate TGF-beta1-induced negative regulation of IL-1R/TLR signaling.” Biochem Biophys Res Commun 393(4): 836–843.

Letterio, J. J. and A. B. Roberts (1998). “Regulation of immune responses by TGF-beta.” Annu Rev Immunol 16: 137–161.

Li, M. O. and R. A. Flavell (2006). “TGF-beta, T-cell tolerance and immunotherapy of autoimmune diseases and cancer.” Expert Rev Clin Immunol 2(2): 257–265.

Martinez, J. D., J. A. C. Garza and A. Cuellar-Barboza (2019). “Going Viral 2019: Zika, Chikungunya, and Dengue.” Dermatol Clin 37(1): 95–105.

Netea, M. G., F. L. van de Veerdonk, J. W. van der Meer, C. A. Dinarello and L. A. Joosten (2015). “Inflammasome-independent regulation of IL-1-family cytokines.” Annu Rev Immunol 33: 49–77.

New, J. and S. M. Thomas (2019). “Autophagy-dependent secretion: mechanism, factors secreted, and disease implications.” Autophagy: 1–12.

Pandey, N., A. Jain, R. K. Garg, R. Kumar, O. P. Agrawal and P. V. Lakshmana Rao (2015). “Serum levels of IL-8, IFNgamma, IL-10, and TGF beta and their gene expression levels in severe and non-severe cases of dengue virus infection.” Arch Virol 160(6): 1463–1475.

Pang, T., M. J. Cardosa and M. G. Guzman (2007). “Of cascades and perfect storms: the immunopathogenesis of dengue haemorrhagic fever-dengue shock syndrome (DHF/DSS).” Immunol Cell Biol 85(1): 43–45.

Patra, G., S. Mallik, B. Saha and S. Mukhopadhyay (2019). “Assessment of chemokine and cytokine signatures in patients with dengue infection: A hospital-based study in Kolkata, India.” Acta Trop 190: 73–79.

Perez, A. B., B. Sierra, G. Garcia, E. Aguirre, N. Babel, M. Alvarez, L. Sanchez, L. Valdes, H. D. Volk and M. G. Guzman (2010). “Tumor necrosis factor-alpha, transforming growth factor-beta1, and interleukin-10 gene polymorphisms: implication in protection or susceptibility to dengue hemorrhagic fever.” Hum Immunol 71(11): 1135–1140.

Sierra, B., A. B. Perez, K. Vogt, G. Garcia, K. Schmolke, E. Aguirre, M. Alvarez, F. Kern, G. Kouri, H. D. Volk and M. G. Guzman (2010). “Secondary heterologous dengue infection risk: Disequilibrium between immune regulation and inflammation?” Cell Immunol 262(2): 134–140.

Soo, K. M., B. Khalid, S. M. Ching, C. L. Tham, R. Basir and H. Y. Chee (2017). “Meta-analysis of biomarkers for severe dengue infections.” PeerJ 5: e3589.

Srikiatkhachorn, A., A. Mathew and A. L. Rothman (2017). “Immune-mediated cytokine storm and its role in severe dengue.” Semin Immunopathol 39(5): 563–574.

St John, A. L. and A. P. S. Rathore (2019). “Adaptive immune responses to primary and secondary dengue virus infections.” Nat Rev Immunol 19(4): 218–230.

Tan, T. Y. and J. J. Chu (2013). “Dengue virus-infected human monocytes trigger late activation of caspase-1, which mediates pro-inflammatory IL-1beta secretion and pyroptosis.” J Gen Virol 94(Pt 10): 2215–2220.

Tillu, H., A. S. Tripathy, P. V. Reshmi and D. Cecilia (2016). “Altered profile of regulatory T cells and associated cytokines in mild and moderate dengue.” Eur J Clin Microbiol Infect Dis 35(3): 453–461.

Travis, M. A. and D. Sheppard (2014). “TGF-beta activation and function in immunity.” Annu Rev Immunol 32: 51–82.

Wilder-Smith, A., E. E. Ooi, O. Horstick and B. Wills (2019). “Dengue.” Lancet 393(10169): 350–363.

Wu, M. F., S. T. Chen, A. H. Yang, W. W. Lin, Y. L. Lin, N. J. Chen, I. S. Tsai, L. Li and S. L. Hsieh (2013). “CLEC5A is critical for dengue virus-induced inflammasome activation in human macrophages.” Blood 121(1): 95–106.

Yacoub, S. and B. Wills (2014). “Predicting outcome from dengue.” BMC Med 12: 147.

Yeo, A. S., N. A. Azhar, W. Yeow, C. C. Talbot, Jr., M. A. Khan, E. M. Shankar, A. Rathakrishnan, Azizan, S. M. Wang, S. K. Lee, M. Y. Fong, R. Manikam and S. Devi Sekaran (2014). “Lack of clinical manifestations in asymptomatic dengue infection is attributed to broad down-regulation and selective up-regulation of host defence response genes.” PLoS One 9(4): e92240.

